# Characterization of extracellular matrix deposited by segmental trabecular meshwork cells

**DOI:** 10.1101/2023.03.11.532242

**Authors:** VijayKrishna Raghunathan, Andrews Nartey, Kamesh Dhamodaran, Hasna Baidouri, Julia A. Staverosky, Kate E Keller, Keith Zientek, Ashok Reddy, Ted Acott, Janice A Vranka

**Affiliations:** Department of Basic Sciences, College of Optometry, University of Houston, Houston, Texas; Ophthalmology and Visual Sciences, Casey Eye Institute, Oregon Health & Science University, Portland, Oregon, United States; Proteomics Shared Resources, Oregon Health & Science University, Portland, Oregon, United States

## Abstract

Biophysical and biochemical attributes of the extracellular matrix are major determinants of cell fate in homeostasis and disease. Ocular hypertension and glaucoma are diseases where the trabecular meshwork tissue responsible for aqueous humor egress becomes stiffer accompanied by changes in its matrisome in a segmental manner with regions of high or low flow. Prior studies demonstrate these alterations in the matrix are dynamic in response to age and pressure changes. The underlying reason for segmentation or differential response to pressure and stiffening are unknown. This is largely due to a lack of appropriate models (*in vitro* or *ex vivo*) to study this phenomena. In this study, we characterize the biomechanical attributes, matrisome, and incidence of crosslinks in the matrix deposited by primary cells isolated from segmental flow regions and when treated with glucocorticosteroid. Data demonstrate that matrix deposited by cells from low flow regions are stiffer and exhibit a greater number of immature and mature crosslinks, and that these are exacerbated in the presence of steroid. We also show a differential response of high or low flow cells to steroid via changes observed in the matrix composition. We conclude that although a mechanistic basis for matrix stiffness was undetermined in this study, it is a viable tool to study cell-matrix interactions and further our understanding of trabecular meshwork pathobiology.

## INTRODUCTION

Glaucoma is an age associated multifactorial and heterogeneous group of diseases resulting in optic neuropathy encompassing loss of retinal ganglion cells, thinning of the retinal nerve fiber layer, cupping of the optic nerve head resulting in irreversible loss of vision. Reducing intraocular pressure (IOP) continues to remain the only modifiable causal risk factor to slow glaucoma progression. Ocular hypertension (OHT) is largely hypothesized to be a result of increased resistance to outflow of aqueous humor via the trabecular meshwork [TM; cells & extracellular matrix (ECM)], inner wall endothelial cells of the Schlemm’s canal, and distal vasculature(Swaminathan et al., 2014). Amongst these, the primary site of resistance is believed to be the cells and ECM of the juxtacanalicular trabecular meshwork (JCT)(Acott et al., 2021). Specifically, accumulation of ECM and plaque-like material in the JCT have been reported in eyes with primary open angle glaucoma (POAG) (Lütjen-Drecoll et al., 1986a; Lütjen-Drecoll et al., 1986b; Rohen et al., 1993; Ueda et al., 2002), and such changes are associated with stiffening of the TM (Last et al., 2011; Vahabikashi et al., 2019). In fact, prior work from our group demonstrates that such stiffening is segmental with regions of high flow (HF) being softer while regions of low flow (LF) are stiffer in glaucoma. This corresponds to loss of a homeostatic response to elevated pressure (Raghunathan et al., 2018).

A growing body of literature documents that HF and LF regions show differential molecular expression of ECM and matricellular proteins and genes suggesting that cells in segmental regions and their matrices may differ intrinsically (Keller et al., 2011; Vranka and Acott, 2017; Vranka et al., 2015). It is now well recognized that cells sense and respond to cues in their extracellular microenvironment differently (Baker and Chen, 2012; Kechagia et al., 2019; Kim et al., 2011; Lu et al., 2011), thus it is likely that TM cell fate decisions to external stressors would influence homeostatic responses to elevated pressures. Prior studies demonstrate that normal human anterior segments perfused at 1x or 2x pressure results in a differential biomechanical response of segmental flow regions accompanied by corresponding changes in the ECM (Vranka et al., 2018). These studies demonstrated that HF regions may serve a compensatory role by becoming softer when LF regions increase in stiffness at elevated pressures. This is comparable with observations in experimental glaucoma where unlasered TM regions of glaucoma non-human primates (NHPs) were comparatively softer than control unlasered NHPs (Raghunathan et al., 2017).

It is currently not well understood why cells in these segmental flow regions respond differently at elevated pressures. We hypothesize that this may be due to multiple factors: **(i)** either cells within segmental flow regions in human TM are distinct and thus sense and respond to the external stressor differently, **(ii)** the extracellular microenvironment of the segmental cells differ sufficiently to enforce differential cellular mechanosensation and mechanotransduction, and/or **(iii)** the ratio and prevalence of various cell types within the segmental regions of the TM differ substantially to elicit differential responses. To the best of our knowledge, while cellular composition of the TM has been investigated from whole TM tissues (Patel et al., 2020; van Zyl et al., 2020), whether differences exist between segmental flow regions remain to be seen. To further our understanding of segmental flow physiology, we recently developed a robust feasible method to isolate, culture, and characterize cells from human segmental flow regions *in vitro* (Staverosky et al., 2020). Further studies highlighted critical differences in the culture conditions, molecular pathways, and responses to effects of biophysical stimuli between various segmental flow cells (Dhamodaran et al., 2022; Staverosky et al., 2020). Here, we expand on these studies and characterize the extracellular matrices deposited by segmental TM cells *in vitro* using atomic force microscopy, immunocytochemistry, and proteomics.

## MATERIALS AND METHODS

### Primary human trabecular meshwork segmental cell isolation and culture

Primary human TM cells (hTM) cells were isolated from glaucomatous or non-glaucomatous donors. Whole globes and/or corneoscleral rims were obtained from eye banks (SavingSight, St. Louis, MO: Lions VisionGift eye bank, Portland, OR). This study is not considered Human Subjects Research because cells were acquired post-mortem from de-identified donor tissues. Therefore, it is deemed exempt by University of Houston’s Institutional Review Board (IRB). Donor demographics and characterization/validation of TM cell strains utilized in this experiment were described and shown previously (Dhamodaran et al., 2022), in accordance with consensus recommendation (Keller et al., 2018), only DEX-responsive cell strains were used in our study. Briefly, the HF and LF regions for cell isolation were identified by perfusion of whole globes with CellMask™ Orange plasma membrane strain (ThermoFisher, Waltham, MA, USA) for 90 min prior to dissection as described previously (Staverosky et al., 2020). Segmental TM tissues were dissected, placed with 0.2% (w/v) collagen coated cytodex beads in growth medium [Dulbecco’s modified Eagle medium/Nutrient Mixture F-12 (50:50; DMEM/F-12) with 2.5 mM L-glutamine supplemented with 10% fetal bovine serum (FBS), and 1% penicillin/streptomycin/amphotericin (Life Technologies, Carlsbad, CA, USA)]. HF and LF cells that migrated out of the segmental TM tissues were maintained in growth media and used between passages two and five for all experiments.

### Derivation of cell derived matrices

Culture of cells and derivation of ECM was performed as described previously (Yemanyi et al., 2020a). Briefly, HF and LF TM cells were cultured in growth medium in the presence or absence of 100 nM dexamethasone (Dex; Sigma-Aldrich Corp., St. Louis, MO, USA) on 3-aminopropyl trimethoxysilane modified glass coverslips in a 24-well tissue culture plate for 4 weeks with media changes/treatment administered every other day. Matrices were obtained by decellularization of cultures as described previously (Yemanyi et al., 2020a) using 20 mM ammonium hydroxide and 0.05% Triton X-100 in deionized water. Freshly decellularized ECM was used for mechanical characterization and proteomics, while formaldehyde-fixed cell-derived matrices were used for immunocytochemistry. Samples treated with vehicle control (ethanol) are referred to as VehM, while those with steroid are referred to as glucocorticoid induced matrices (GIM) henceforth.

### Atomic force microscopy (AFM) for mechanical characterization of cell derived ECM

Elastic moduli of ECM were determined as described previously (Chang et al., 2014; Raghunathan et al., 2018; Yemanyi et al., 2020a; Yemanyi et al., 2020b). Briefly, force-distance curves were obtained in contact mode using PNP-TR cantilevers (spring constant in the range of 0.17-0.24 N/m; Nano and More, USA) with a pyramidal tip modified with 5-10 μm diameter borosilicate beads. For all samples, force curves were obtained from at least 5–10 locations with 5 force curves at each location. All experiments were performed for matrices derived from three individual donor strains. Elastic moduli were determined using Hertzian model for linear elastic materials; details of the equation and analysis are described elsewhere (Chang et al., 2014; Hunter, 1960; Willis, 1966). Thickness of the CDM were determined by a ‘scratch and indent’ method using pyramidal AFM cantilevers and measured between 7-13 μm in thickness.

### Immunocytochemistry

All ECM samples were fixed in 4% paraformaldehyde for 20 min, washed three times each with 0.01% Triton X-100 in phosphate buffered saline (PBS, pH 7.4) and PBS, blocked with 3% fish gelatin, and immunolabelled with anti-collagen IV (catalog # ab6311, Abcam, Cambridge, MA, USA), anti-laminin (catalog # ab11575, Abcam, Cambridge, MA, USA) or anti-Fibronectin (catalog # ab6584, Abcam, Cambridge, MA, USA) followed by species appropriate fluorophore-tagged secondary antibody. Where appropriate, samples were stained with F-actin (catalog # 00045, CF594 Phalloidin, Biotium, Fremont, CA, USA) to confirm decellularization, and nuclei were counterstained with DAPI (catalog # D1306, FisherScientific, CA, USA). All experiments were performed for all cell strains, with at least 3 samples per cell strain. For each immunolabelled sample, 3–8 random locations were imaged using a Leica DMi8 inverted fluorescence microscope. Average fluorescence intensity of immunolabelled samples were quantified and semi-quantitative analyses is shown here. As appropriate, representative images from a single field of view for a single donor are illustrated.

### Proteomic analysis of cell derived matrices

Composition of the ECMs deposited by both glaucomatous and non-glaucomatous cells was determined by shotgun proteomics as described previously (Raghunathan et al., 2015a; Raghunathan et al., 2015b).

#### LC-MS Proteomics Analysis

Cell strain samples were centrifuged, and the pellet was transferred to a clean tube. Cells were suspended in 5% SDS, 50 mM triethylammonium bicarbonate (TEAB) buffer and homogenized using a Sonic Dismembrator 60 ultrasonic probe (Thermo Fisher Scientific, Waltham, MA, USA). Proteins were reduced with dithiothreitol (DTT) and alkylated with iodoacetamide (IAA). Proteins were diluted with 90%/10% methanol/100mM TEAB and bound to an S-Trap column (Protifi, Farmingdale, NY, USA). The S-Trap columns were washed with 90%/10% methanol/100mM TEAB to remove SDS and bound proteins were digested using trypsin. Peptides were eluted from the S-trap column, dried by centrifugal evaporation, and reconstituted in 5% formic acid in water.

Peptides were analyzed by LC-MS on a Dionex Ultimate HPLC coupled to an Orbitrap QE Orbitrap mass spectrometer (Thermo Fisher Scientific, Waltham, MA, USA) operating in Data Dependent Acquisition (DDA) mode. Data was processed using the PAW pipeline (Wilmarth et al., 2009) with the Comet search engine (version 2016.03) (Eng et al., 2013) searching versus the Uniprot proteome UP000005640 (Homo Sapiens, taxon ID 9606) canonical FASTA sequences (81,837 proteins). Searches were conducted with hydroxylation on proline and methionine as a variable modification and carbidomethyl on cysteine as a static modification.

#### LC-MS Crosslink Analysis

Cell strain samples were centrifuged, and the pellet was transferred to a clean tube. Cells were suspended in 1mM NaOH and homogenized using a Sonic Dismembrator 60 ultrasonic probe (Thermo Fisher Scientific, Waltham, MA, USA). Solid NaBH4 was added at a ratio of 1:100 by weight and the samples were allowed to incubate for one hour at room temperature in a well-ventilated fume hood. After incubation the samples were washed three times with ultrapure water. Samples were hydrolyzed by the addition of 6N HCl followed by incubation under UHP nitrogen at 95°C for 18 hours. Hydrolyzed samples were dried by centrifugal evaporation and reconstituted in 0.2% heptaflourobutyric acid (HFBA) in water.

Crosslinks were analyzed by LC-MS on an Agilent 1100 HPLC coupled to a Velos Pro mass spectrometer (Thermo Fisher Scientific, Waltham, MA, USA) operating in Parallel Reaction Monitoring (PRM) mode. DPD and PYD standards were used to confirm the retention times of the analytes analyzed in the samples. Data was analyzed and peak areas were calculated using the Skyline software package (Pino et al., 2020).

### Statistical analysis

#### AFM

For ECM mechanics, force-distance curves from the AFM were exported as an ASCII text file into MATLAB programming, and a custom program was used to determine the elastic modulus of cell-derived ECM for the groups. Using repeated measures ANOVA, we tested the hypothesis that the mean elastic modulus of HF VehM, LF VehM, HF GIM, and LF GIM are the same using a significance level of 0.05. Consequently, a posthoc Tukey test was used to make pairwise comparisons.

#### Immunocytochemistry

Although fluorescent images of cell derived ECM, like other fluorescent images are typically qualitative, we sought to compute quantitative values to aid comparison of our cell derived ECM groups. We thus used an in-built feature called mean fluorescence on our immunofluorescent microscope to provide quantitative values for the ECM that were expressed by our samples. All statistical analyses were performed using R Programming language. For n=3 biological cell strains, 4 samples per strain, and three technical replicates per sample, we captured 4 fields of view for each replicate. We performed a one-way ANOVA to test the hypothesis that there is no difference in the mean quantitative cell derived ECM deposited by our four samples: HF cells treated with vehicle (HF VehM), LF cells treated with vehicle (LF VehM), HF cells treated with DEX (HF GIM), LF cells treated with DEX (LF GIM) at a significance level of 0.05. We proceeded to perform post hoc Tukey test to compare all paired groups.

#### Gene Ontology using the Panther Classification System

Proteins identified from proteomics analyses were classified by the Gene Ontology PANTHER classification system (Mi and Thomas, 2009). Gene Ontology is defined as the framework for the model of biology and classifies functions along three aspects: molecular function (molecular activities of gene products), cellular component (where gene products are active), and biologic process (pathways and larger processes made up of the activities of multiple gene products (details can be found at: http://geneontology.org/ and http://www.pantherdb.org/ in the public domain).

## RESULTS

Prior studies have demonstrated differences in the ECM/matricellular genes between segmental flow regions, and that the tissue resident cells respond differently to pressure challenge *ex vivo* (Acott et al., 2020; Keller et al., 2011; Raghunathan et al., 2018; Vranka et al., 2015; Vranka et al., 2020; Vranka et al., 2018). Recently we developed a method to isolate cells from segmental flow regions and demonstrated that they may intrinsically differ suggesting their potential utility for mechanistic studies (Staverosky et al., 2020). Here, we investigate biochemical and biomechanical differences in ECM deposited by these cells *in vitro*, in the absence or presence of dexamethasone, and characterize them further.

First, we determined the expression of three ECM proteins that are components of the TM outflow resistance: collagen IV, laminin, and fibronectin (**Figure 1**). Since we were interested in only ECM that was deposited, we did not quantify differences in secreted proteins. In the absence of Dex, no significant differences in the morphology or expression levels (semi-quantitative) of fibronectin and collagen IV were observed comparing HF and LF cells. A small, yet significant difference was observed for laminin. In the presence of Dex, the relative fluorescence intensity of all three ECM proteins were greater than that observed without Dex. Further, a subtle but significant difference was observed comparing HF *vs* LF for all 3 ECM proteins when cells were treated with Dex.

**Figure 1:**
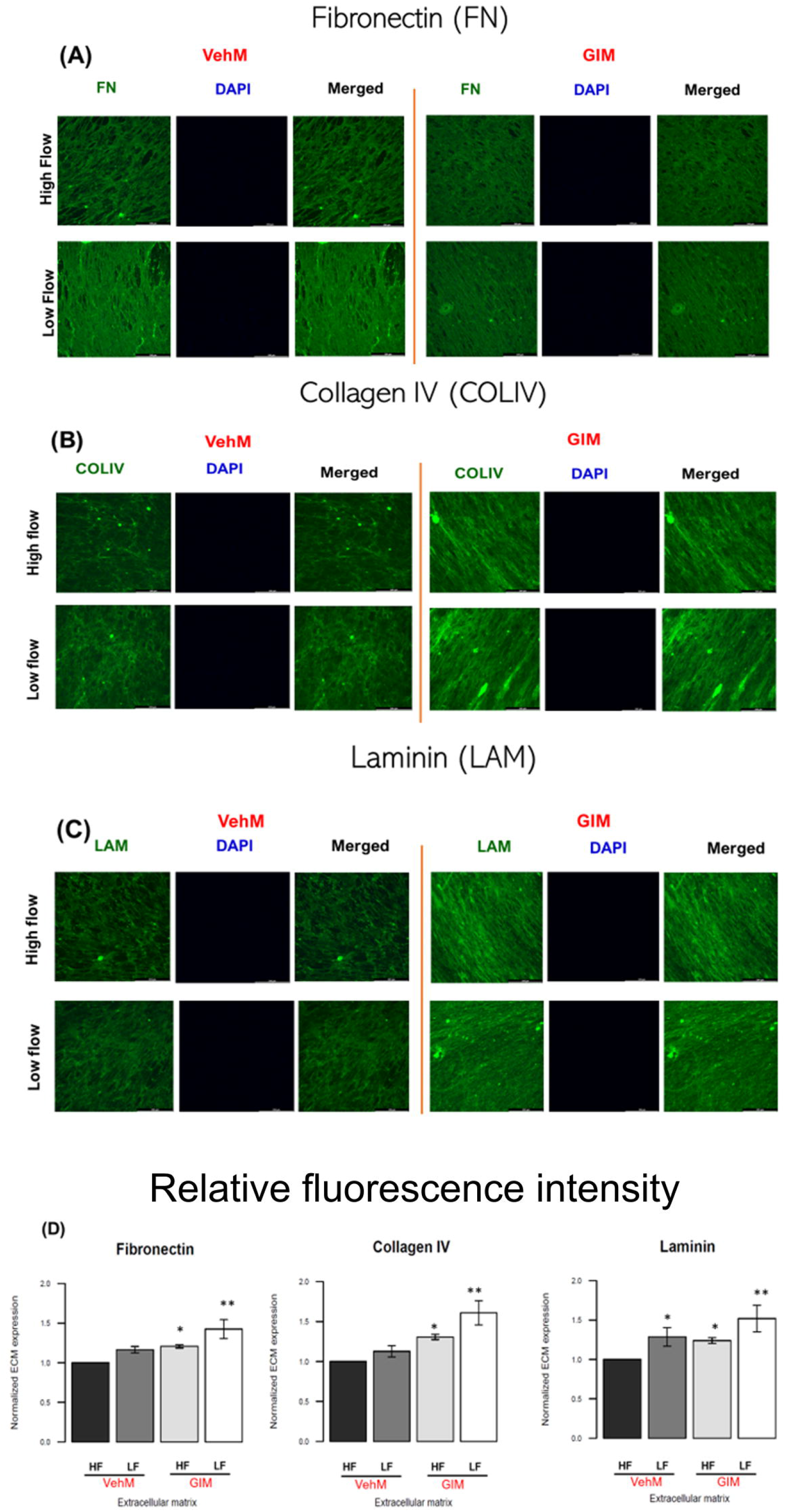
Differential expression patterns are seen comparing ECM [**(A)** Fibronectin, **(B)** Collagen IV, **(C)** Laminin] from HF and LF cells in the presence or absence of dexamethasone. Images are representative of those observed from each group. **(D)** Relative fluorescence intensity was quantified from at least 4-6 fields of view for each donor/group, and a minimum of 3 donors were used for these experiments. Bar graphs are mean ± standard deviation (n=3 donors). *p<0.05, **p<0.01 compared with HF VehM group, one-way ANOVA, followed by Tukey’s multiple comparison test.

Next, we sought to quantify the elastic moduli of ECM deposited by HF *vs* LF cells by atomic force microscopy (**Figure 2**). The mean elastic moduli of the samples were: HF VehM: 0.737 ± 0.161 kPa (n = 3 biological replicates); LF VehM: 1.391 ± 0.264 kPa (n = 3 biological replicates); HF GIM: 2.205 ± 0.209 kPa (n = 3 biological replicates); LF GIM: 3.668 ± 0.855 kPa (n = 3 biological replicates). Statistically significant differences were observed in moduli values between HF and LF ECM both in the presence and absence of Dex suggesting intrinsic differences between the matrix deposited by the cells, and their response to steroid treatment.

**Figure 2:**
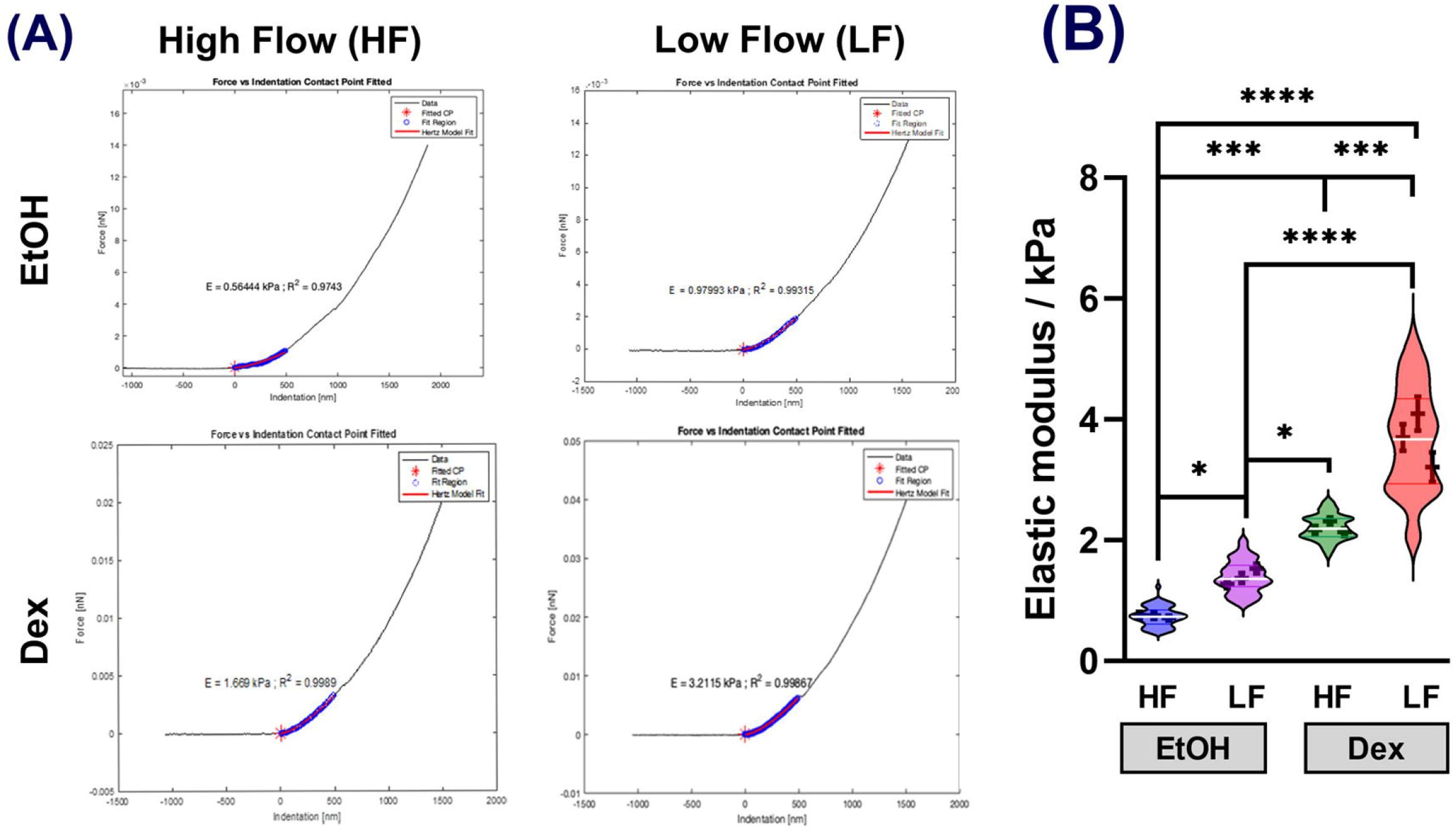
Biomechanical characterization of cell derived matrices from HF and LF cells in the presence/absence of dexamethasone. **(A)** Graphs illustrate representative force-indentation curves with Hertzian fit. **(B)** Violin plot of elastic moduli measures from each donor/group from 9-10 individual locations for each sample. Mean ± standard error in mean are indicated in the error bars for individual donors. *p<0.05, **p<0.01, ***p<0.001, ****p<0.0001, two-way ANOVA, followed by Tukey’s multiple comparison test.

Subsequently, we sought to investigate the factors resulting in ECM moduli differences. One factor that has been attributed to mediate tissue/matrix mechanics is the prevalence of crosslinks between matrix proteins. Using mass spectrometry, we quantified the presence of immature (dihydroxylysinonorleucine; DHLNL) and mature [deoxy-pyridinoline (DPD) and pyridinoline (PYD)] collagen crosslinks in the ECM deposited by HF and LF cells in the presence or absence of Dex (**Figure 3**). In the absence of Dex, we observed that the ECM deposited by LF cells yielded a higher amount of immature crosslinks (DHLNL) compared with those by HF cells. Due to large variability, statistical significance was unattained. In the presence of Dex, no statistically significant differences in the peak areas for DHLNL were observed when comparing HF and LF cells. Further, no significant differences in DHLNL content in the CDMs were observed comparing control *vs* steroid treated cells (**Figure 3A**). In quantifying mature crosslinks, we fist investigated the ratio of PYD:DPD which is an indicator of lysyl hydroxylase (LH1) activity and collagen stabilization (**Figure 3B**). We observed that the ratio of PYD:DPD was greater in ECM deposited by LF cells compared with HF cells, although statistical significance was only observed between LF VehM (LF EtOH) and HF GIM (HF Dex) groups. To determine the contribution of PYD to total mature crosslinks, we determined the ratio of PYD:(PYD+DPD) and observed no significant differences (**Figure 3C**). Further, no differences were observed comparing the ratios of immature to mature crosslinks in the CDMs between segmental cells regardless of Dex treatment (**Figure 3D**). Finally, by performing a linear regression between elastic modulus of the CDMs *vs* the crosslink maturity ratio [defined as ratios of immature to mature crosslinks i.e. DHLNL:(PYD+DPD)], we demonstrated no relationship between elastic modulus and presence of crosslinks (**Figure 3E**).

**Figure 3:**
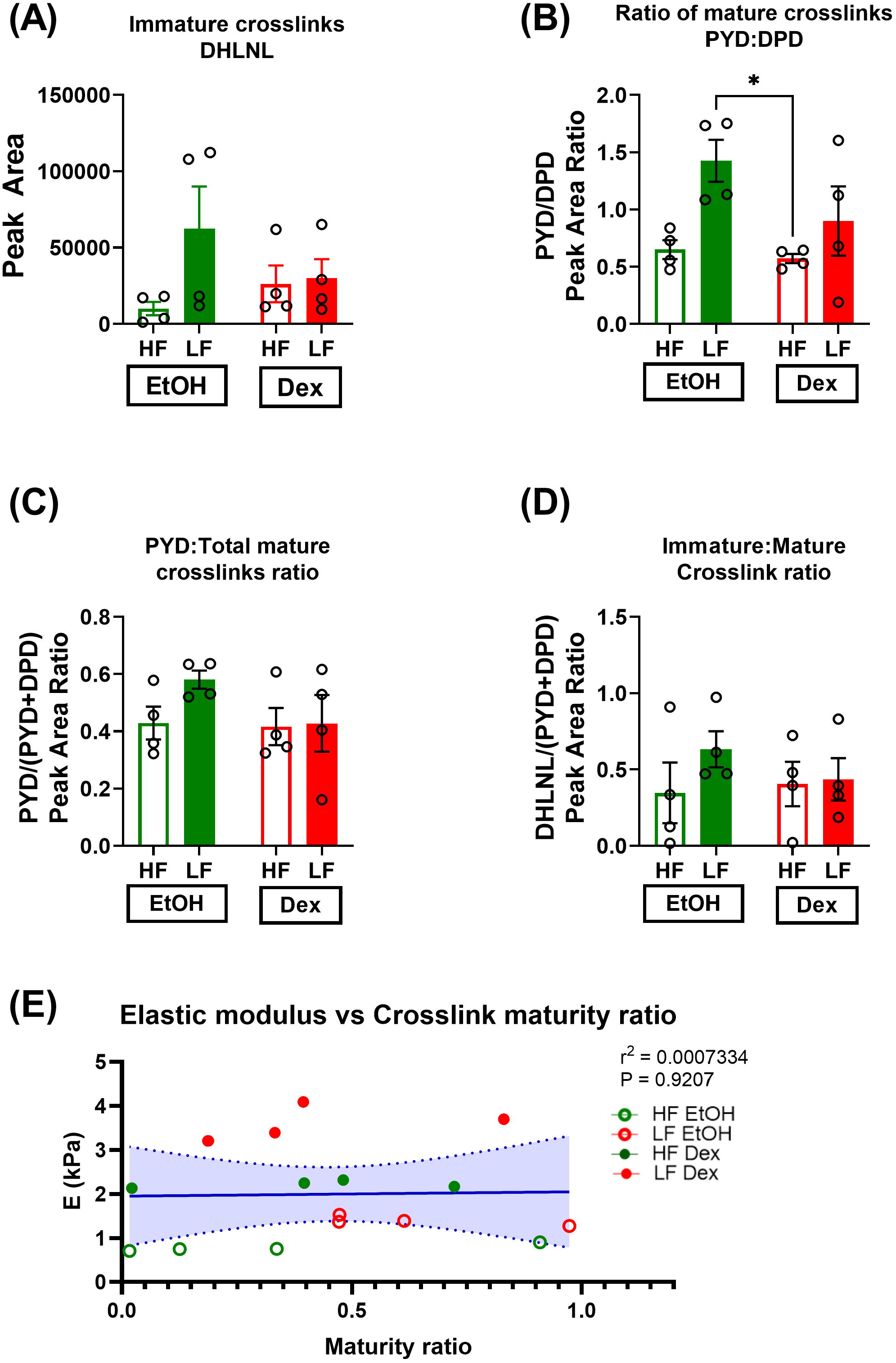
Quantification of mature and immature collagen crosslinks in cell derived matrices HF and LF cells in the presence/absence of dexamethasone. Values were determined from area of chromatographic peaks obtained by LC/MS. Graphs are **(A)** immature crosslinks (DHLNL), **(B)** ratio of mature crosslinks (PYD:DPD), **(C)** ratio of PYD to total mature crosslinks [i.e. PYD: (PYD+DPD)], **(D)** Crosslink maturity ratio - defined as ratio of immature to total mature crosslinks [i.e. DHLNL:(PYD+DPD)], **(E)** correlation between elastic modulus and crosslink maturity ratio. Dotted lines represent 95% confidence interval. Histograms represent mean ± standard error in mean (n=4 donors), *p<0.05, two-way ANOVA, followed by Tukey’s multiple comparison test.

Next, we performed quantitative proteomics to determine changes in protein expression in the ECM deposited by HF and LF cells in the presence and absence of dexamethasone. Proteins altered 2-fold in expression consistently in 3 to 4 cell lines were only considered in our analysis. First, comparing HF *vs* LF ECM from control (VehM/EtOH) group, we observed 33 proteins upregulated while 43 were downregulated in ECM of LF cells (**Figure 4A**). Comparing Dex treated HF *vs* LF groups (GIM), a significant number of proteins (95 proteins) were downregulated in the LF group while 3 proteins were upregulated (**Figure 4B**). Next, we observed that the ECM deposited by HF cells yielded 23 proteins upregulated with 25 proteins downregulated after Dex treatment (**Figure 4C**). In contrast, ECM deposited by LF cells yielded 3 proteins upregulated with 97 proteins downregulated after Dex treatment (**Figure 4D**). Using the STRING database (Szklarczyk et al., 2019) the aforementioned protein list was then hierarchically clustered and mapped into a functional protein-protein association network for ECM derived from HF and LF cells in the presence or absence of Dex (**Figure 5A-D**).

**Figure 4:**
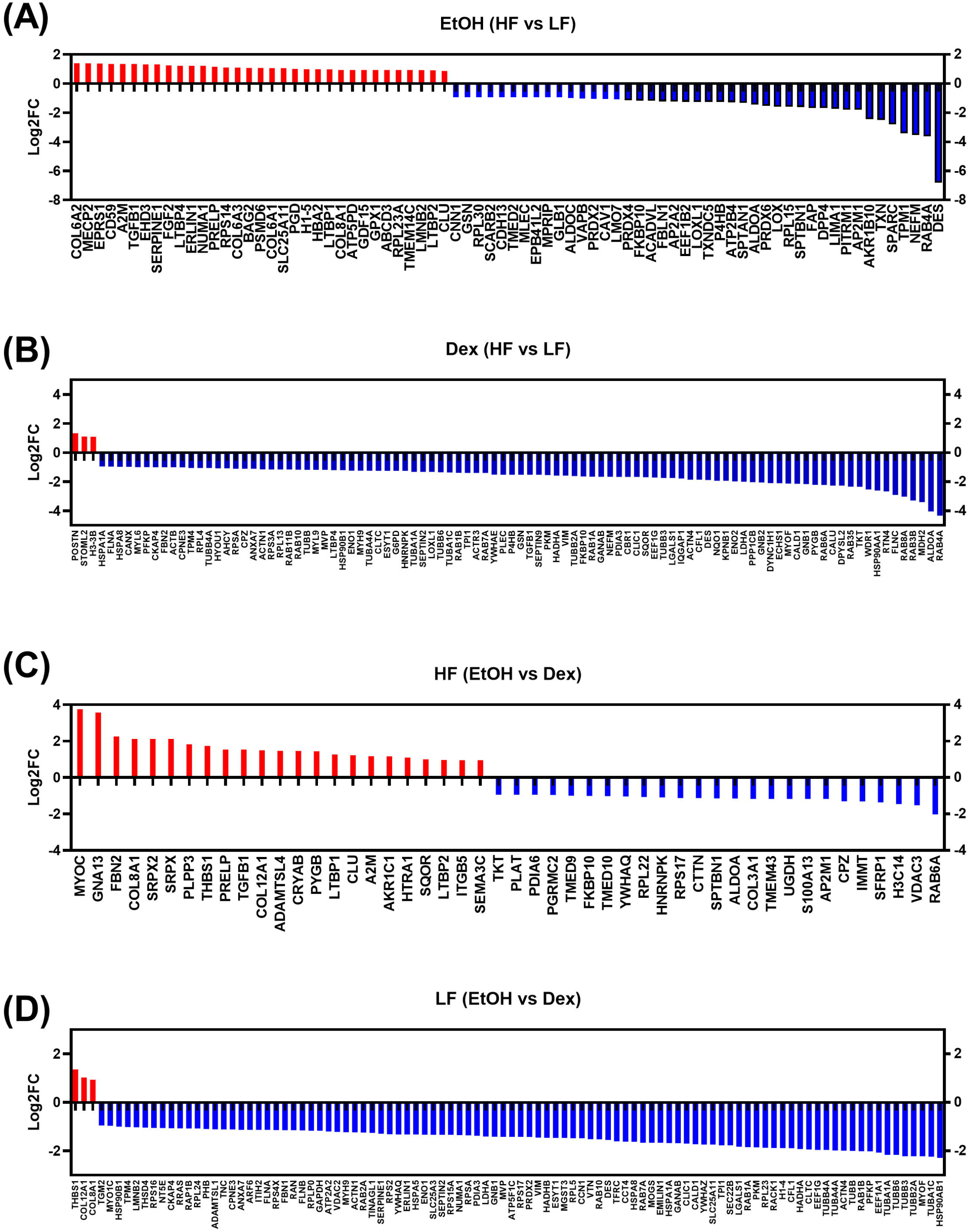
Histograms illustrating log to the base 2, two-fold changes of protein expression in ECM derived from segmental flow cells in the presence/absence of dexamethasone. Proteins that were upregulated are in red and those downregulated are in blue. Graphs represent comparisons of proteins identified in ECM derived from **(A)** LF and HF cells in the absence of any treatment, **(B)** LF and HF cells in the presence of 4 weeks of dexamethasone treatment, **(C)** HF cells in the presence or absence of 4 weeks of dexamethasone treatment, **(D)** LF cells in the presence or absence of 4 weeks of dexamethasone treatment.

**Figure 5:**
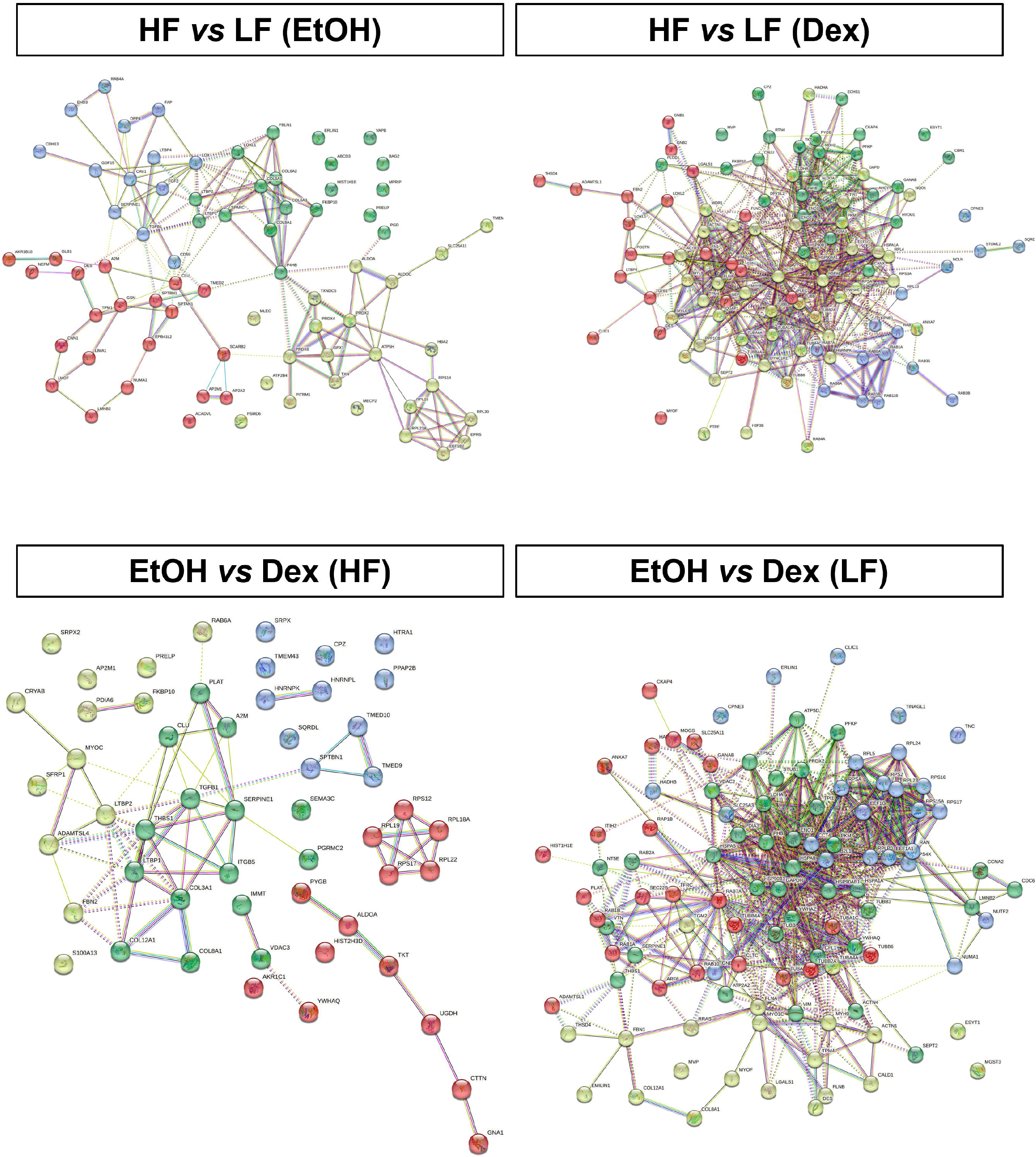
Protein-protein interaction maps of two-fold changes of protein expression in ECM derived from segmental flow cells in the presence/absence of dexamethasone.

Subsequently, the differentially expressed protein candidates were classified using the Gene Ontology PANTHER classification system for biological processes (**Figure 6**) and protein class (**Figure 7**). In LF VehM ECM compared with HF VehM ECM (EtOH group), 34% of the upregulated proteins and 32% of the downregulated proteins represent proteins that regulate cellular processes. The major upregulated proteins were extracellular matrix and metabolite interconversion proteins, while oxidoreductases represented the majority of downregulated proteins. In LF GIM ECM compared with HF GIM ECM (Dex group), 40% of the upregulated and 36% of the downregulated proteins represent proteins that regulate cellular processes. Classes of proteins that are enriched in either group were diverse and with none particularly overrepresented in LF GIM ECM compared with HF GIM ECM.

**Figure 6:**
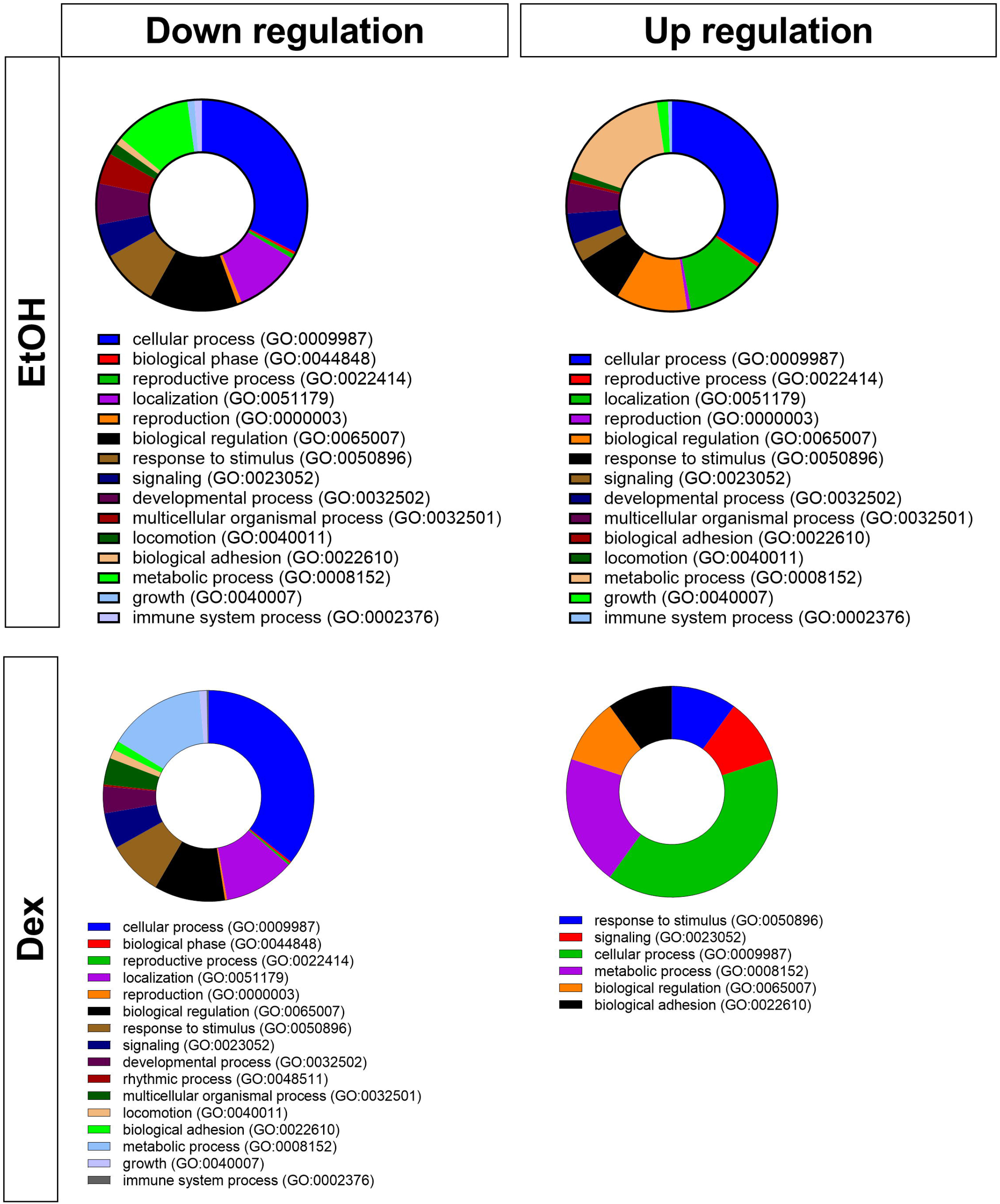
Gene ontology classification of biological processes, represented in pie charts as a percentage of the total number of proteins identified, comparing ECM from HF *vs* LF cells either in the presence or absence of dexamethasone.

**Figure 7:**
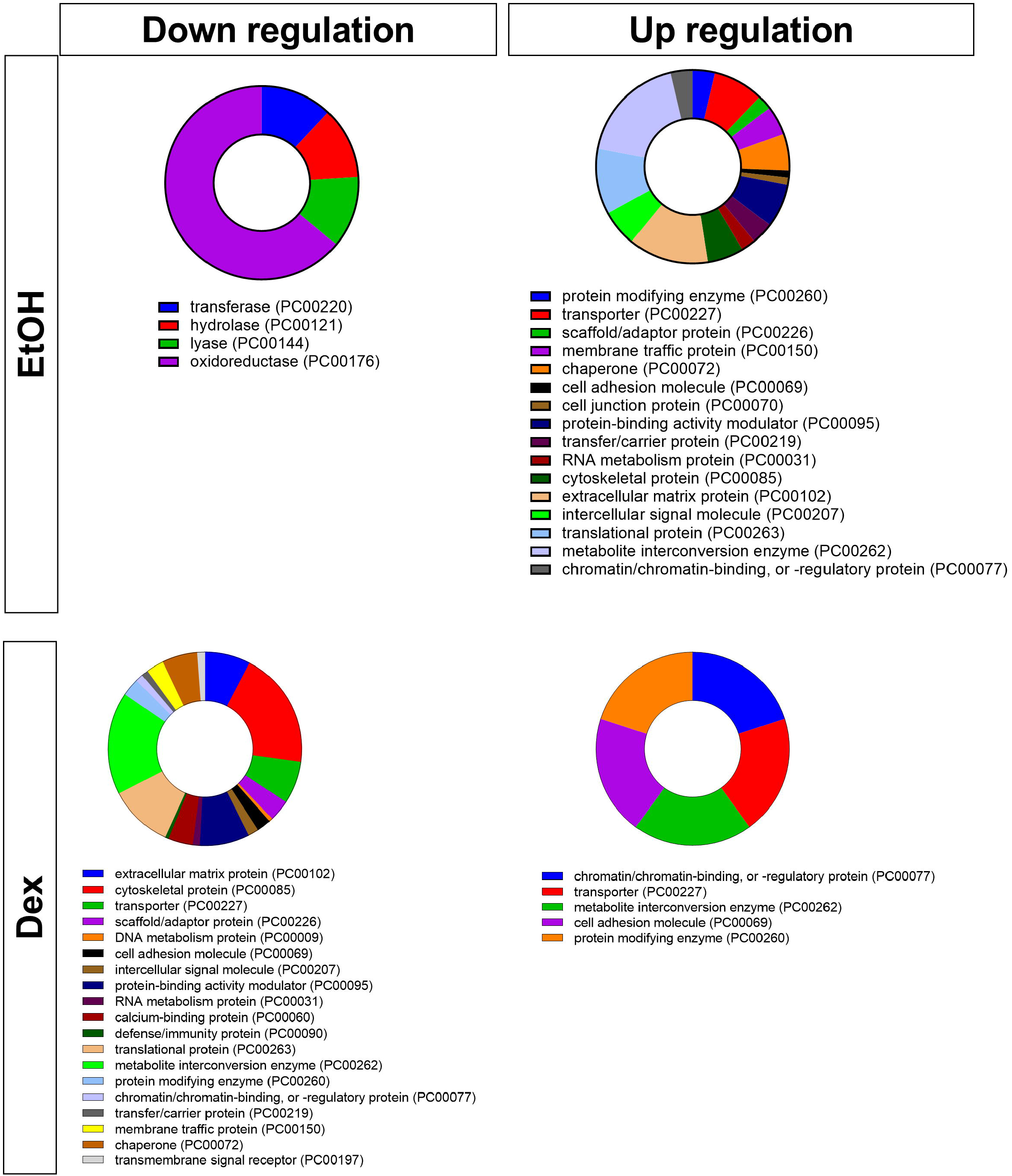
Gene ontology classification of protein class, represented in pie charts as a percentage of the total number of proteins identified, comparing ECM from HF *vs* LF cells either in the presence or absence of dexamethasone.

## DISCUSSION

Outflow of aqueous humor outflow across the TM is non-uniform and segmental. Recent studies from our group and others have demonstrated that such segmentation is dynamic in the absence of external stimulus in live rodents *in vivo* (Reina-Torres et al., 2022) and with elevated pressure in *ex vivo* organ perfusion models (Vranka et al., 2020). These physiological changes are associated and partly mediated through regional differences in biomechanical and biochemical composition of HF and LF tissues (Vranka et al., 2018) (Keller et al., 2011; Vranka and Acott, 2017; Vranka et al., 2015). Since differences in ECM are observed locally, we posited that distinct cellular identity in these localized segments likely contribute to these phenomena. Indeed, intrinsic differences in expression levels of select genes, some matricellular, were reportedly observed between segmental flow cells during early characterization and validation *in vitro* (Dhamodaran et al., 2022; Staverosky et al., 2020). Building on these prior observations, we were motivated to further characterize the ECM deposited by primary hTM cells isolated and cultured from segmental flow regions and in response to chronic steroid treatment *in vitro*.

Semi-quantitative analyses of key ECM proteins (Fibronectin, Collagen IV, and Laminin), typically present in the TM (Acott and Kelley, 2008; Acott et al., 2021), show an upward trend in their expression patterns in the absence of any stimulus. Dexamethasone stimulation resulted in the overexpression of these proteins in matrix both HF and LF cells, although the response was strongest in LF cells. Steroid induced overexpression of critical matrix proteins is consistent with well-established TM cell culture studies and in animal models (Clark and Wordinger, 2009; Liesenborghs et al., 2020; Overby et al., 2014; Raghunathan et al., 2015b; Shinzato et al., 2007; Zhou et al., 1998). Response of segmental TM tissue to steroid treatment in *ex vivo* anterior segment perfusion systems is yet to be comprehensively evaluated, although changes in total TM biomechanics (cells, matrix, and tissue) have been reported previously (Peng et al., 2022; Raghunathan et al., 2015b). Using cell derived matrices as a model, Dex induced stiffening of the ECM has also been shown *in vitro* (Raghunathan et al., 2015b). Here, we show that regardless of Dex treatment, ECM deposited by LF cells was stiffer than that deposited by HF cells. Chronic steroid treatment exacerbated this phenotype.

After mechanical characterization, we sought to investigate whether the relative distribution of ECM proteins differed across the groups. Indeed, we observed differences in subsets of proteins in ECM deposited by HF *vs* LF cells. In the absence of Dex, ECM of LF cells (LF VehM) was over-represented with key matricellular and proteins such as COL6A2/3, COL8A1, CLU, GDF15, LTBP1/2/4, TGFβ1, and SERPINE1. Most of these matrix proteins have garnered special attention by the community previously. For example, GDF15 has been implicated with visual field loss in pseudo exfoliation glaucoma and with increased contractility and ECM production in the TM (Lin et al., 2020; Muralidharan et al., 2016). More recently, a rodent model with a total loss of clusterin was shown to be associated with OHT *in vivo*, while clusterin supplementation *in vitro* attenuated the pro-fibrotic effects of TGFβ2 (Soundararajan et al., 2022). In its extracellular secreted form, clusterin works together with the plasminogen activation system to facilitate the clearance of extracellular misfolded proteins and their aggregates (Constantinescu et al., 2017; Satapathy and Wilson, 2021). Along with SERPINE1 (Plasminogen activator inhibitor-1), clusterin is a secreted marker for hypoxia-inducible factor (Nakamura et al., 2006). Constitutive upregulation of LTBP and TGFβ are both associated with activation of the TGFβ pathway and suggestive of an increased association with fibronectin extradomain A (FN-EDA) that have all been long implicated in TM dysfunction (Annes et al., 2003; Bollinger et al., 2011; Danford et al., 2017; Fleenor et al., 2006; Fuchshofer et al., 2003; Klingberg et al., 2018; Munoz et al., 2006; Saharinen et al., 1999). In the presence of Dex, a number of proteins that mediate cellular response to ECM changes, cytoskeletal remodeling, and cell adhesion are differentially modulated. Collectively, these suggest that the ECM deposited by LF cells differ substantially from that deposited by Dex both in the presence and absence of Dex, and that these matrices may be primed with factors that activate stress responsive and TGFß pathways.

While ECM accumulation and ultrastructure have been extensively studied in TM tissues (Hann et al., 2001; Johnson et al., 1997; Rohen et al., 1993), exacting their role *in vitro* has been more challenging due to technical limitations in model development. Indeed, even though several changes in protein expression levels are observed in this study, these by themselves do not explain the increased stiffness of the matrix. Changes in tissue / matrix biomechanics have often attributed to either **(i)** ECM accumulation, **(ii)** ECM morphology/ultrastructure, or **(iii)** ECM crosslinking. In order to determine if crosslinking of ECM was a factor in reduced outflow facility, it was demonstrated that overexpression of tissue transglutaminase (TGM2), a potent crosslinker of collagen, resulted in OHT while its knockdown impaired the effects of TGFß2 induced OHT (Raychaudhuri et al., 2017; Raychaudhuri et al., 2018). Recent efforts to understand the role of, and type of, crosslinks in disease have gained attention. Typically, collagen crosslinks take two forms: mature and immature, which largely depends on the process by which they are formed. Interfribrillar and intrafibrillar crosslinking of collagen fibers may be formed by both enzymatic (e.g., via lysyloxidases (LOX)) and non-enzymatic (e.g. via glycation processes (AGEs)) processes that determine the structural integrity and mechanical properties of the ECM (Avery et al., 2009; Eyre et al., 1991). Enzymatic crosslinking is relatively quicker and yields mature crosslinks (Moriguchi and Fujimoto, 1978; Takahashi et al., 1995). The ratio of immature to mature crosslinks is a good indicator of the amount of matrix remodeling (Berteau et al., 2015) i.e. a greater ratio is suggestive of fewer mature crosslinks and thus more remodeling. Our data here demonstrate that, though the number of immature and mature crosslinks themselves trended higher in LF ECM compared with HF, the ratio of immature to mature crosslinks was comparable regardless of Dex treatment. The increasing trend observed in the number of crosslinks suggests it may play a partial role in determining ECM stiffness, but this alone may be insufficient to explain the correlation to mechanical properties.

It is thought that changes to the matrix in disease or with age occur over several years *in vivo*. It is thus feasible that 4 weeks of incubation period *in vitro* may be insufficient to truly elicit chronic crosslinking in the ECM. That said, a recent manuscript showed that supplementation of LOXL2 (an isoform of LOX) aided the formation of collagen fibrils (within 2 weeks) /fibers (within 4 weeks) / fascicles (within 6 weeks) in a temporal manner (Bates et al., 2023). Specifically, the authors demonstrated that the timing of LOXL2 addition was critical in determining increases in mature PYD accumulation and alterations to tissue mechanics. In this study, we did not document temporal changes in crosslinking enzymes, their isoforms, or their activity levels. Nevertheless, at least *in vitro*, interpreting our data for ECM remodeling remains inconclusive under the experimental conditions of this study. Further studies are required to ascertain the type of crosslinks in TM tissue and what the causal factors may be. Finally, that subtle differences in matrix deposition, composition and mechanics exist between HF and LF cells suggests that there may be some element of mechanical memory retained in these cells post-isolation/culture, and may be consequential for mechanotransduction events in TM physiology. The epigenetic or molecular mechanisms that underlie mechanical memory in cells (Lele et al., 2020; Mathur et al., 2020; Peng et al., 2017) are beyond the scope of this study.

## CONCLUSION

To the best of our knowledge, this is the first study to comprehensively investigate if there are innate differences in the matrix that primary hTM cells from segmental flow regions deposit. Here we demonstrate that individual differences in matrix stiffness, and ECM protein expression exists but that the causal factor for ECM stiffening may not be explained by crosslinks alone. Nevertheless, we provide a proof-of-concept that protein crosslinks may be determined from cell derived matrices. Further studies are essential to determine how ECM accumulation, ultrastructural changes, and crosslinks work in tandem to elicit changes in matrix biomechanics. Importantly, our data demonstrates that cells isolated and propagated from segmental flow regions may be a viable tool to study cell-ECM interactions and further our understanding of TM pathobiology.

## ACKNOWLEDGEMENT

We thank the Lions VisionGift and SavingSight (Kansas City, MO, USA) for procuring all human donor eyes used in this work. Most importantly, we would like to thank the families of the organ donors without whose consent these experiments would be impossible. We pay our gratitude and respects to Dr. Janice A Vranka, who passed away in August 2021. She was a dedicated scientist, colleague, mentor, and was passionate about science and furthering our understanding of segmental trabecular meshwork biology. Her contributions to this study are enormous and greatly appreciated.

